# Ontogenetic Color Switching in Lizards as a by-Product of Guanine Cell Development

**DOI:** 10.1101/2022.01.12.475993

**Authors:** Gan Zhang, Venkata Jayasurya Yallapragada, Michal Shemesh, Avital Wagner, Alexander Upcher, Iddo Pinkas, Harry L.O. McClelland, Dror Hawlena, Benjamin A. Palmer

## Abstract

Many animals undergo dramatic changes in colour during development^1,2^. Changes in predation risk during ontogeny are associated with spectacular switches in defensive colours, typically involving the replacement of skin or the production of new pigment cells^3^. Ontogenetic colour systems are ideal models for understanding the evolution and formation mechanisms of animal colour which remain largely enigmatic^2^. We show that defensive colour switching in lizards arises by reorganization of a single photonic system, as an incidental by-product of chromatophore maturation. The defensive blue tail colour of hatchling *A. beershebensis* lizards is produced by light scattering from premature guanine crystals in underdeveloped iridophore cells. Camouflaged adult tail colours emerge upon reorganization of the guanine crystals into a photonic reflector during chromatophore maturation. The substituent guanine crystals form by the attachment of individual nanoscopic plates, which coalesce during growth to form single crystals. Our results show that the blue colour of hatchlings is a fortuitous, but necessary, precursor to the development of adult colour. Striking functional colours in animals can thus arise not as distinct evolutionary innovations but *via* exploitation of the timing of naturally occurring changes in chromatophore cell development.

## Main

Many animals undergo ontogenetic colour changes driven by variations in habitat, diet, size and predation^3-5^. Changes in predation risk during ontogeny induce spectacular switches in defensive colours from masquerade to aposematic^6^ or from cryptic to less-cryptic^7^, involving integument replacement or pigment cell production. A classic example of colour switching is found in hatchling lizards, whose active foraging renders crypsis ineffective. Hatchling lizards use colourful, autotomizable tails to redirect predator attacks away from the vital organs to increase survival probability^8-13^. The conspicuous tail colours fade during ontogeny as the animals adopt less risky behaviours^8,11-15^. Key questions are: How do transiently functional defensive colours evolve? What are the optical mechanisms and physiological costs underlying ontogenetic colour changes^16^? How do the component photonic structures form^17^?

We elucidate the ontogenetic colour change mechanism in the tail of the Be’er Sheva fringe-fingered lizard (*Acanthodactylus beershebensis*). The blue tail colour of hatchlings is produced by light-scattering from premature guanine crystals in underdeveloped iridophore cells. White/brown adult tail colours emerge upon reorganization of the guanine crystals into a photonic reflector during chromatophore maturation. The defensive blue colours of hatchlings are thus a consequence of delayed chromatophore development (i.e., a type of colour neoteny), which enables them to actively forage with lower mortality risk. This lizard model shows that guanine photonic reflectors, which are widely distributed in animals^18^, can form *via* gradual re-orientational ordering of crystals from disordered precursor states. Guanine crystals form by the attachment of individual nanoscopic platelets, which independently nucleate and coalesce during growth to form single coherent crystals - reminiscent of oriented attachment growth mechanisms^19^.

### Ontogenetic changes in lizard tail colours

We followed colour changes in *A. beershebensis* lizards during ontogeny. The lizards hatch with a conspicuous, matte blue tail, whose reflectance spectrum exhibits a single broad peak at ∼460 nm (Fig. 1a, d). Within two weeks, the blue tail colour fades, and a small peak at ∼550 nm emerges - broadening the reflectivity spectrum (Fig. 1b, d). By adulthood, the tail transforms into a cryptic white/brown colour (Fig. 1c) exhibiting broadband reflectivity, with minor peaks at ∼550 nm and ∼630 nm (Fig. 1d).

**Fig. 1.**
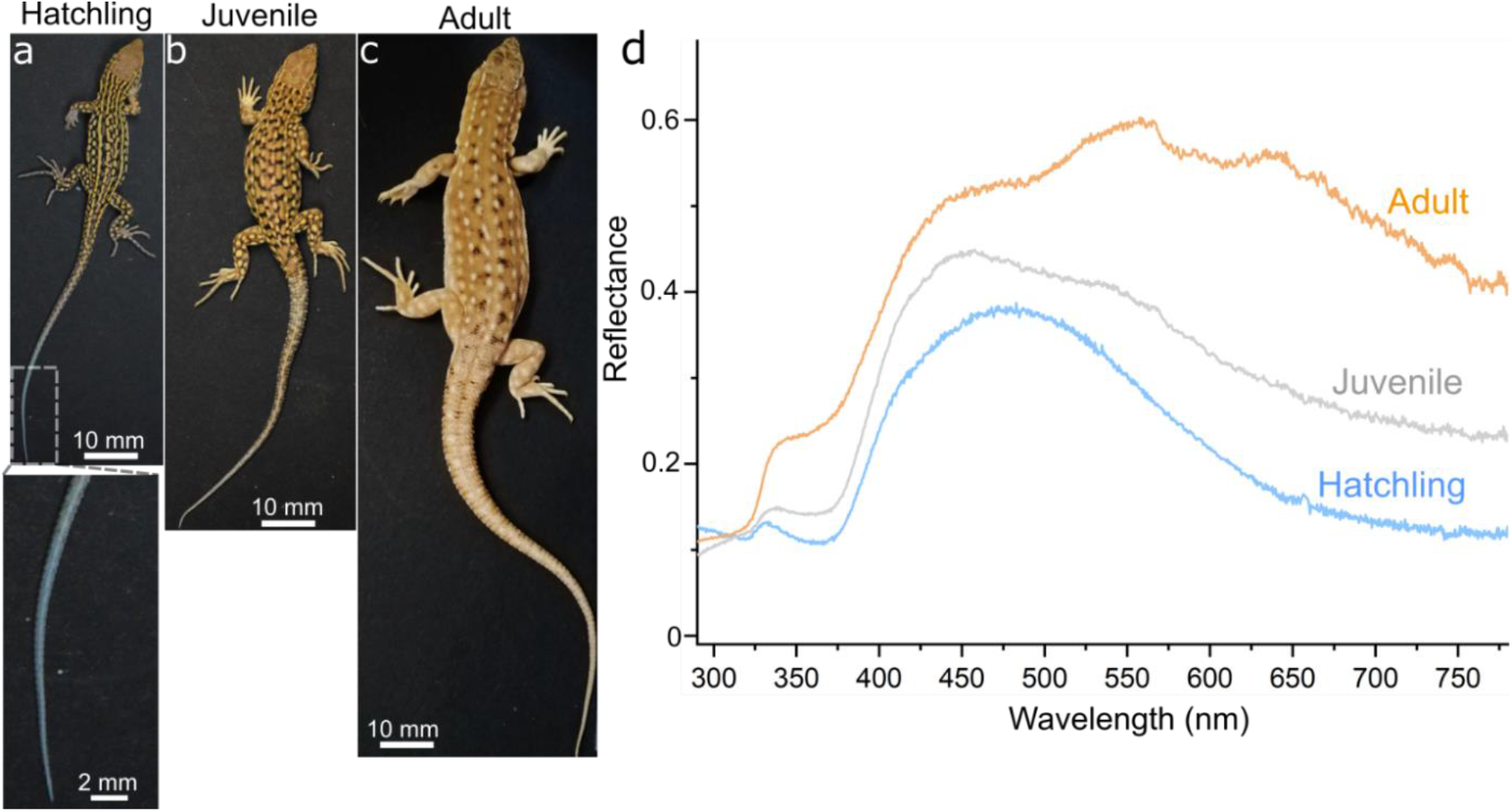
Optical properties of *A. Beershebensis* tails during ontogeny. (**a** to **c**) Dorsal-lateral views of hatchling (inset; higher magnification view of the hatchling’s blue tail), juvenile and adult *A. Beershebensis*. **(d)** Reflectance spectra from posterior regions of the tails in different developmental stages. The total reflectance increases with development. All spectra exhibit a small peak in the UV at ∼340 nm and a sharp drop in reflectance intensity from ∼450 to 375 nm.

### Chromatophore maturation during ontogeny

Skin colour in lizards is produced by layers of different pigment cells, collectively termed - the dermal chromatophore unit^20^. To elucidate the mechanism underlying this colour change, we imaged the ultrastructure of the component chromatophore cells during ontogeny. The uppermost xanthophores contain pteridine and carotenoid pigments and function as spectral filters^21-24^. Guanine crystals in the underlying iridophores produce reflective colours due to their high refractive index and nanostructural organization^18,25-28^. Light transmitted through the guanine layer is absorbed by the lowest melanophore layer^20,22^. Polarizing optical micrographs of cross-sections through hatchling (Fig. 2a, b) and adult tails (Fig. 2f, g) reveal highly birefringent, dermal iridophores, which, in adults, are interspersed with xanthophores. TEM micrographs (Fig. 2c, d) reveal that hatchling xanthophores contain empty pigment granules (pterinosomes)^29^, exhibiting electron-dense inclusions and membrane invaginations (Fig. 2d, black and white arrows) – features of premature pterinosomes^30^. A few weeks after hatching (Fig. 2h), pigment condenses inside the pterinosomes, which develop an onion-like lamellae structure (Fig. 2i, j, red arrow) – characteristic of mature pterinosomes^30,31^. In both hatchlings and adults, the iridophores contain angular void spaces – indicative of guanine crystals lost during sample preparation for TEM (Fig. 2e, i)^32-34^. The number of iridophore layers in a transverse section of the dermis increases from 2 in hatchlings to 4-5 in adults.

**Fig. 2.**
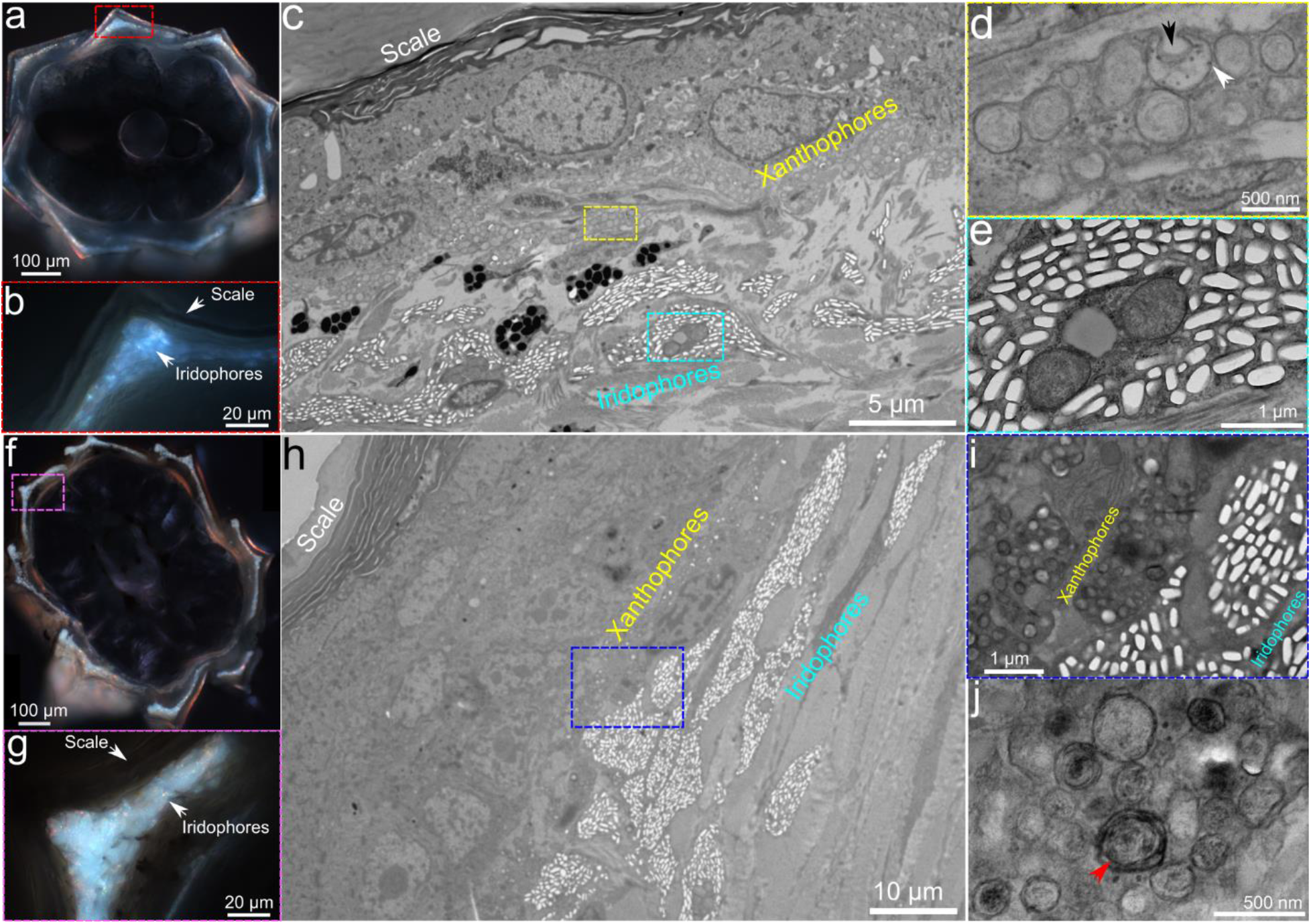
Optical and TEM micrographs of cross-sections through hatchling and adult tails. **(a, b)** Optical micrographs of a hatchling tail. (**c**) Low magnification TEM image of a hatchling tail. (**d, e**) High magnification TEM images of hatchling xanthophores and iridophores. (**f, g**) Optical micrographs of an adult tail. (**h**) Low magnification TEM image of an adult tail. (**i, j**) High magnification TEM images of adult xanthophores and iridophores.

We used cryogenic-scanning electron microscopy (cryo-SEM) to image the xanthophores and iridophores at different development stages in their native, hydrated state (Extended Data Fig. 1). Hatchling iridophores contained randomly oriented, membrane-bound, ellipsoidal vesicles (Fig. 3a,) together with a few faceted crystals. Fractures through these vesicles reveal minute, partially formed guanine crystals (Fig. 3b, Extended Data Fig. 2) constructed from separate nanoscopic platelets (platelet thickness: ∼12 nm) and surrounded by aqueous material^32,33^. Vesicles containing the smallest crystals had a circular cross-section (Fig. 3c, blue arrow). More mature crystals extend across the width of the vesicle, causing the membrane to re-shape around the growing crystal, adopting its morphology (i.e., an oval cross-section, Fig. 3c, red arrow, Extended Data Fig. 2). Similarly, hatchling xanthophores are also underdeveloped (Fig. 3d, e). Pterinosome granules are filled by smoothly textured, aqueous material and often exhibit membrane invaginations as observed in TEM (Fig. 3d, cyan arrow). Cryo-SEM images of juveniles (∼ 2 weeks post-hatching) show the transition of the chromatophores towards maturation. More faceted vesicles are observed in the iridophores as the membranes condense around the surface of the forming crystals (Fig. 3f). At this transitional stage of ontogeny the crystals, though still orientationally disordered, begin to exhibit a preferred orientation. The distinctly separated platelets of the immature hatchling crystals have coalesced to form single, coherent guanine crystals (Fig. 3g, h, Extended Data Fig. 2, 3, 4)^35,36^. At this stage, pterinosome granules in the xanthophores exhibit a more fibrous character (Fig. 3i, j)^30^.

**Fig. 3.**
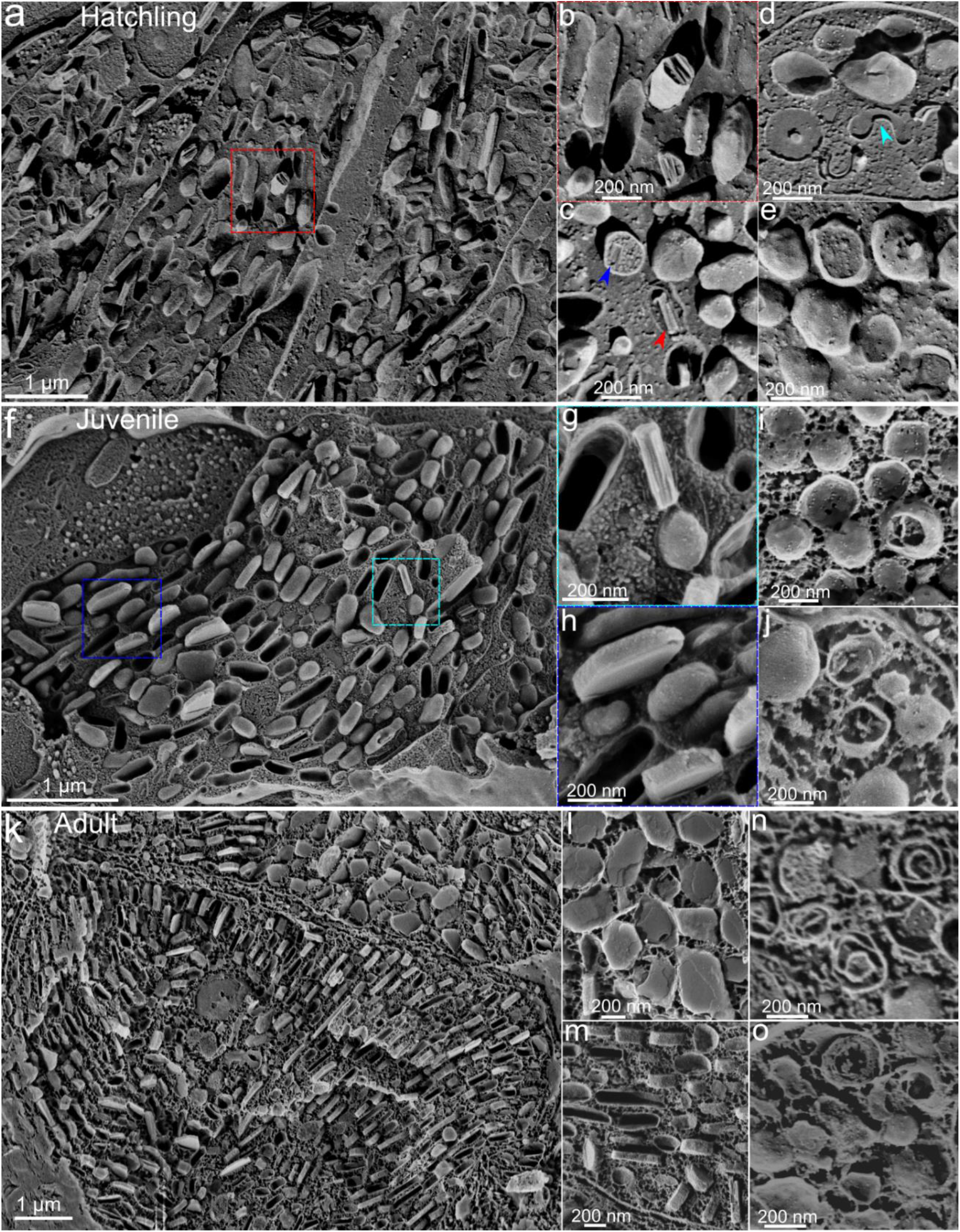
Cryo-SEM micrographs of the chromatophores during ontogeny. **(a)** Clusters of iridophores in a hatchling tail. **(b, c)** High magnification images of the crystal-vesicles in the iridophores. Minute, partially formed crystals composed of nanoscopic crystal layers are observed in the fractured vesicles (**c** is taken from a different region of the sample). **(d, e)** High magnification images of fractured pigment granules in the hatchling xanthophores. **(f)** A single iridophore in the juvenile stage. (**g, h**) High magnification images of guanine crystals from the same iridophore. **(i, j)** High magnification images of pterinosome vesicles containing fibrous structures in juvenile xanthophores. **(k)** Iridophores in an adult tail packed with mature guanine crystals. (**l, m**) High magnification images of the mature guanine crystals oriented in different directions. (**n, o**) Mature pterinosomes with concentric lamellar structures in the adult xanthophores.

In the adult, the iridophores are larger than in the hatchling and juveniles and are filled with mature, irregular polygonal crystals (mean size ∼ 500 × 270 × 85 nm^3^). Few ellipsoidal vesicles are observed. The crystals are arranged into locally ordered domains containing stacks of crystals separated by cytoplasm – a multilayer reflector (Fig. 3k). Each oriented domain occupies an area of approximately 3×3 μm^2^. Individual domains in a single iridophore are misoriented with respect to one another (Fig. 3 k, l, m). Pigment granules in the xanthophores exhibit an onion-like fibrous structure reminiscent of mature pterinosomes (Fig. 3n, o)^30^.

### Modelling the optical properties of the iridophores

Based on a knowledge of the crystal size and organization obtained from cryo-SEM, we performed electromagnetic calculations to rationalize how the optical properties of the iridophore cells change during ontogeny. In hatchlings, long-range orientational and positional ordering between crystals are absent. Therefore, we expect the optical properties of such a structure to primarily result from the scattering response of individual particles. Finite-difference time-domain (FDTD) calculations^37,38^ (Fig. 4a) showed that the total scattering cross sections exhibit higher scattering in the blue region of the spectrum, which can account for the blue colour of the hatchling tail. However, the shape of the spectrum, which has a peak in the blue (∼ 470 nm) and a drop in reflectance at shorter wavelengths suggests the presence of some structural correlations such as layering (Extended Data Fig. 5).

**Fig. 4.**
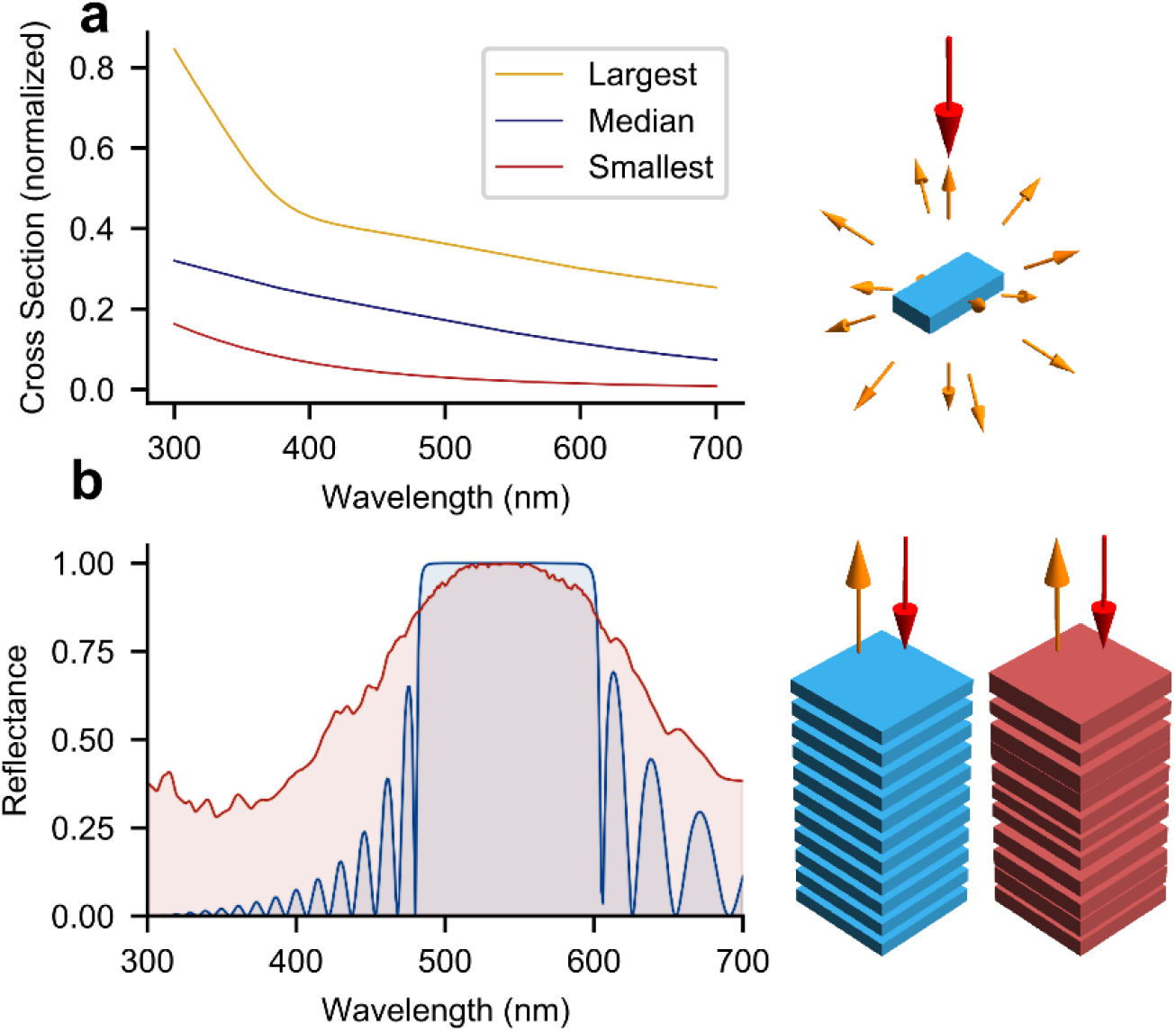
Calculated scattering and reflectivity properties of the iridophores during development. **(a)** Calculated scattering spectra of single guanine crystals of various crystal sizes observed in hatchling iridophores. The spectra exhibit higher scattering in the blue region of the spectrum. Blue trace: median crystal size (325 nm × 185 nm × 50 nm); red trace: smallest crystals: 150 nm × 85 nm × 35 nm; yellow trace: largest crystals (500 nm × 270 nm × 85 nm). The spectra are the average for two orthogonal polarizations. **(b)** Calculated reflectance spectra of multilayer stacks of guanine with crystal thicknesses and cytoplasm spacings derived from cryo-SEM of adults. Blue trace: reflectance spectrum of a periodic stack of 20 layers of guanine (crystal thickness; 81 nm, cytoplasm spacing; 88 nm, estimated values from a single oriented domain). Red trace: Average reflectance spectrum upon introducing disorder in crystal thicknesses and spacings (using standard deviation of crystal thickness and spacings derived from measurements of a single oriented domain). The periodic and disordered stacks are depicted as blue and red stacks in the schematics.

The guanine crystals in the iridophores of the adult are larger, with a length of ∼ 500 nm (mean crystal thickness: 85 ± 25 nm) and are clearly organized into stacks. We modelled the optical properties of such a structure as a multilayer reflector whose constituent layers exhibit disorder in crystal thickness and spacing between crystals. The values of crystal thickness, cytoplasm spacing, and disorder were estimated empirically from cryo-SEM data (Extended Data Fig. 5). The calculated reflectance of a stack of 20 evenly spaced guanine/cytoplasm layers exhibits a high-reflectance plateau centered around 500 nm, which is a clear signature of a photonic bandgap (Fig. 4b, blue curve)^39^. We modelled disordered stacks by introducing random deviations from mean values of crystal thickness and cytoplasm spacing. The magnitude of the disorder observed experimentally by cryo-SEM is sufficient to broaden the reflectance band significantly (Fig. 4b, red curve). In any specific area of the skin, the experimental reflectance spectrum (Fig. 1d) is an average over several such disordered stacks, each characterized by a specific broadened reflectance spectrum, resulting in the broad plateaus observed in the experimentally measured spectra (Fig. 1d). Our models account for the major qualitative features of the spectra that arise from the organization of guanine crystals in iridophores. While structural correlations in the iridophore photonic structures lead to lower reflectance in the deep blue and UV, scattering and absorption from other constituents of the measured skin samples, such as collagen fibers and pigment also affect the spectrum^27^.

## Discussion

Most blue colours produced by guanine-based systems, including those of brightly-coloured lizard tails in other taxa^13,40^ are generated by highly ordered multilayers^22,40-42^. Constructive interference from these Bragg reflectors results in highly saturated, iridescent colours^18,43^. The observation of angle-independent, scattering blue colours from guanine has not been shown before, though Bagnara suggests that some frog mutants appear blue due to incoherent scattering from iridophores^41^. The iridophores in the tails of *A. beershebensis* transform from blue scatterers to broad-band photonic reflectors during ontogeny.

Essentially nothing is known about the formation of photonic structures in living skin (i.e., involving pigment cells)^17^, including widely distributed guanine systems^18^. However, work on butterflies^44-46^, beetles^47,48^, and birds^49,50^ show that photonic systems made primarily of dead tissue (e.g., cuticle, feathers) form by self-assembly processes. In this study, we observe the transformation of a disordered guanine scatterer into a partially ordered photonic device *via* the gradual orientational ordering of crystals (Fig. 3). This transformation is correlated with an increase in density and size of crystals whose plate morphology favours assembly into a multilayer superstructure. This provides a clue as to how architecturally complex photonic structures can be formed from simpler, disordered states, which may also be optically functional at all stages of formation. The constituent guanine crystals form by the attachment of individual nanoscopic platelets, which coalesce during growth to form single, coherent crystals (Fig. 3, Extended Data Fig. 2). This indicates that each individual guanine crystal has multiple nucleation points but that the individual platelets are lattice-matched which is reminiscent of oriented attachment growth mechanisms^19^. The growing guanine crystals effectively re-shape their surrounding membrane, which condenses on the growing surfaces of the crystal, adopting its faceted morphology. Thus, in this case, guanine crystal morphology is not dictated by a pre-shaped template as has been seen in many other biominerals^51^.

Knowledge of this formation process also yields new insights into the ecology and evolution of defensive colours in animals. Many lizards hatch with conspicuous blue tails that serve to deflect predator attacks away from vital organs. In most species, the blue colour fades to concealing colours as lizards grow^8,11-15^, raising the question of how this transiently functional defense strategy evolved? TEM and cryo-SEM imaging, together with optical calculations, show that, in the absence of xanthophore pigmentation^26,33,52^, the blue tail colour of hatchlings arises from diffuse scattering by premature iridophores. Unlike most lizards, snakes, and amphibians, which hatch with fully developed chromatophores, *A. beershebensis* emerges from its egg with premature tail chromatophores. The functional blue tail colour is thus a consequence of delayed chromatophore cell development. Similar observations in *A. boskianus* and *A. scuttelatus* (Extended Data Fig. 6) indicate that this may be a common phenomenon in this genus, and, perhaps in other matte blue-tailed lizards, undergoing ontogenetic colour change. Iridescent blue colours observed in other lizard taxa, which are produced by a different, multilayer interference mechanism, suggests a convergent evolutionary scenario. This scenario is supported by a phylogenetic analysis showing that blue tails evolved 25 times in lizards (unpublished results).

Canonically, ontogenetic colour changes were thought to necessitate the replacement of one optimized optical structure with another (e.g., via integument replacement or the production of new pigment cells). However, our data show that the functional blue colour of hatchling lizards is not an optimized target of evolution but a necessary precursor to the development of adult colours. Colour switching then arises as a by-product of chromatophore cell maturation without the development of a novel, expensive colour-producing mechanism. This raises the question of why selection has favoured delayed chromatophore development and transient blue tail coloration in hatchling lizards. We hypothesize that natal dispersal, competition for limited resources and potentially other ecological factors (e.g., prey size and availability) favours active foraging in hatchlings^53^. Active foraging renders crypsis less effective and selects for alternative defensive strategies and coloration. Lizards with delayed chromatophore development may have a selective advantage because they hatch when the tail is still blue, which increases their probability of surviving inevitable attacks by predator. As the chromatophores develop and the blue colour is lost, lizards must become less active to reduce detectability. To our best knowledge, this is the first example of partial trait neoteny associated with defensive coloration.

Elucidating the ontogenetic colour-change mechanism in this lizard has shown how functional colours in animals can evolve by exploitation of naturally occurring changes in chromatophore cell development. Moreover, this system provides important insights into the assembly of guanine-based photonic structures and the crystallization mechanism of the substituent guanine crystals (by a layered growth mechanism). It remains to be seen whether this is a general phenomenon in all guanine models.

## Methods

### Specimen Collection and Preparation

All specimens of *A. beershebensis* at different stages (hatchling, juvenile and adult) were collected in the Loess Park Nature Reserve, Northern Negev, Israel (permit number 2020/42592). The live lizards were calmed down in the refrigerator at 4 °C for 10 mins for optical microscopy and the reflectance measurements. The last 1 cm of the lizard tails were trimmed and fixed in 4% paraformaldehyde in 1 × PBS fixative in dark environment. The tails were fixed for more than 72 hours and stored in refrigerator at 4 °C until analysed. The fixed tails were placed in 1 × PBS solution for 2 hours and then embedded in 7% agarose gel (1 × PBS as the medium) and cut into 180 μm horizontal sections by a vibratome (LEICA VT1000S). All lizards were released at the exact location of capture in less than 4 hours.

### Optical microscopy

The optical microscopy and polarized light microscopy imaging on fixed vibratome sectioned samples was performed using a Zeiss AX10 microscope equipped with a Zeiss Axiocam 705 colour camera using reflection, transmission or polarization mode with 5×, 10×, 20× and 50× air objectives.

### Reflectance measurements

The tails were cut from the live lizards and then directly measured by an Ocean Optics DH-2000 light source (200-2500 nm) and an Ocean Insight FLAME miniature Spectrometer. The light of the source was injected in one channel of a THORLABS RP20 Reflection probe and the light reflected on the lizard tails ample (near normal incidence) was collected by the second optical fibre channel to the detector. A white diffuse reflectance standard (Labsphere USRS-99-010, AS-01158-060) was used as reference.

### Cryo-SEM imaging

The fresh cut vibratome sections were directly sandwiched between two metal discs (3 mm in diameter, 100 μm in depth for each) and then cryoimmobilized in a high-pressure freezing device (LEICA ICE, High Pressure Freezer). The frozen samples were transferred into a freeze-fracture device (LEICA ACE900 Freeze-Fracture vacuum chamber) by a vacuum cryotransfer sample holder (LEICA VCT500 Cryo-stage) under liquid nitrogen. The samples were freeze-fractured and transferred to a High-resolution GEMINI-300 Zeiss Scanning Electron Microscope. During SEM imaging, the Cryo-stage sample holder and the electron microscope were maintained at -120 °C by liquid nitrogen.

### TEM imaging on fixed and stained Ultra-thin biological tissue

The cut lizard tails were fixed in a fixation solution of 4% paraformaldehyde, 2% glutaraldehyde in 0.1M Cacodylate buffer at pH 7.4 ^27, 40^. The tails were fixed in the buffer for 12-16 hours at room temperature and then rinsed 4 times in cacodylate buffer, then post-fixed and stained with 1% osmium tetroxide, 1.5% potassium ferricyanide in 0.1M cacodylate buffer for 1 hour. Tissues were then washed 4 times in cacodylate, dehydrated in increasing concentrations of ethanol, rinsed twice in propylene oxide, infiltrated with increasing concentrations of Agar 100 resin in propylene oxide, and embedded in fresh resin. Ultra-thin (80 nm) cross-sections were cut with a diamond knife on a LKB 3 microtome. The cross-sections were then placed on carbon-formvar coated copper grids and were post-stained with uranyl acetate and lead citrate. TEM imaging on the ultra-thin biological tissues was performed at a Tecnai 12 TEM 100kV (Phillips) equipped with MegaView II CCD camera.

### TEM imaging and electron diffraction on reflecting materials

The reflecting materials were isolated from the 4% paraformaldehyde fixed lizard tail tissues and cleaned in purified water, and a drop of the resulting suspension was placed on a glow-discharged Cu meshed TEM grid and allowed to dry. The resulting samples were observed with a FEI Tecnai T12 G^2^ TWIN TEM operating at 120 kV. Images and electron diffraction patterns were recorded using a Gatan 794 MultiScan CCD camera.

### *In situ* Micro Raman spectroscopy

The fixed lizard tails and the vibratome sections were placed on an Al metal disc, and the measurements were made using a LabRAM HR Evolution confocal system (Horiba France) with 4 laser lines (325, 532, 632 and 785 nm) and a modular microscope (BX-FM Olympus) with a 1024×256-pixel front illuminated open electrode cooled CCD camera. The confocal microscope had different objectives (LMPlanFL N 50×NA=0.5, MPlanFL N 100×NA=0.9, and MPlanFL N 150×NA=0.9 were used as needed), which allowed accurate spectra acquisition (with a spatial resolution of about 1 μm at 100× objective).

### Optical modelling

The scattering cross section of individual guanine crystals were calculated using the finite-difference time-domain (FDTD) technique, by integrating over the field scattered by the particle illuminated by a linearly polarized plane wave. The particle was oriented such that the face with the largest area is normal to the direction of propagation of incident light. Reported cross sections are the average of values obtained with two orthogonal polarizations. Simulations were performed using a commercial grade electromagnetic solver ^38^. Reflectance of multilayer stacks has been calculated using a 4×4 transfer matrix program. Disorder was introduced in the form of random increments to spacing and thickness, drawn from Gaussian distributions whose width is determined from the variance estimated from Cryo-SEM images. The spectra are an average of a hundred trial spectra. In all calculations the refractive index of guanine was taken to be n_o_ = 1.86, n_e_ = 1.46 and the spacing around/between the particles is assumed to possess a refractive index of 1.33.

## Acknowledgements

We thank Profs. Devi Stuart-Fox, Yael Lubin and Yehu Moran for scientific discussions and advice on the manuscript. This work was supported by an ERC Starting Grant (‘CRYSTALEYES’) and an Azrieli Foundation Faculty Fellowship awarded to B.A.P, and by Gans Collections and Charitable Fund donation to D.H. B.A.P. is the Nahum Guzik Presidential Recruit. G.Z. is the recipient of the Kreitman Postdoctoral Fellowship. Electron microscopy studies were supported by Ilse Katz Institute for Nanoscale Science & Technology at Ben-Gurion University of the Negev.

## Author contributions

B.A.P., G.Z., and D.H. designed the study. G.Z. and A.W. performed cryo-SEM measurements. M.S. and D.H. provided lizard specimens and TEM data. V.J.Y. carried out the scattering and reflectivity calculations. A.U. and I.P. performed guanine crystal characterization. B.A.P., G.Z., D.H. and H.L.O.M. wrote the manuscript with contributions from all the authors.

## Competing interests

The authors declare no competing interests.

**Extended Data** is available for this paper.

## Extended Data

**Extended Data Fig. 1.**
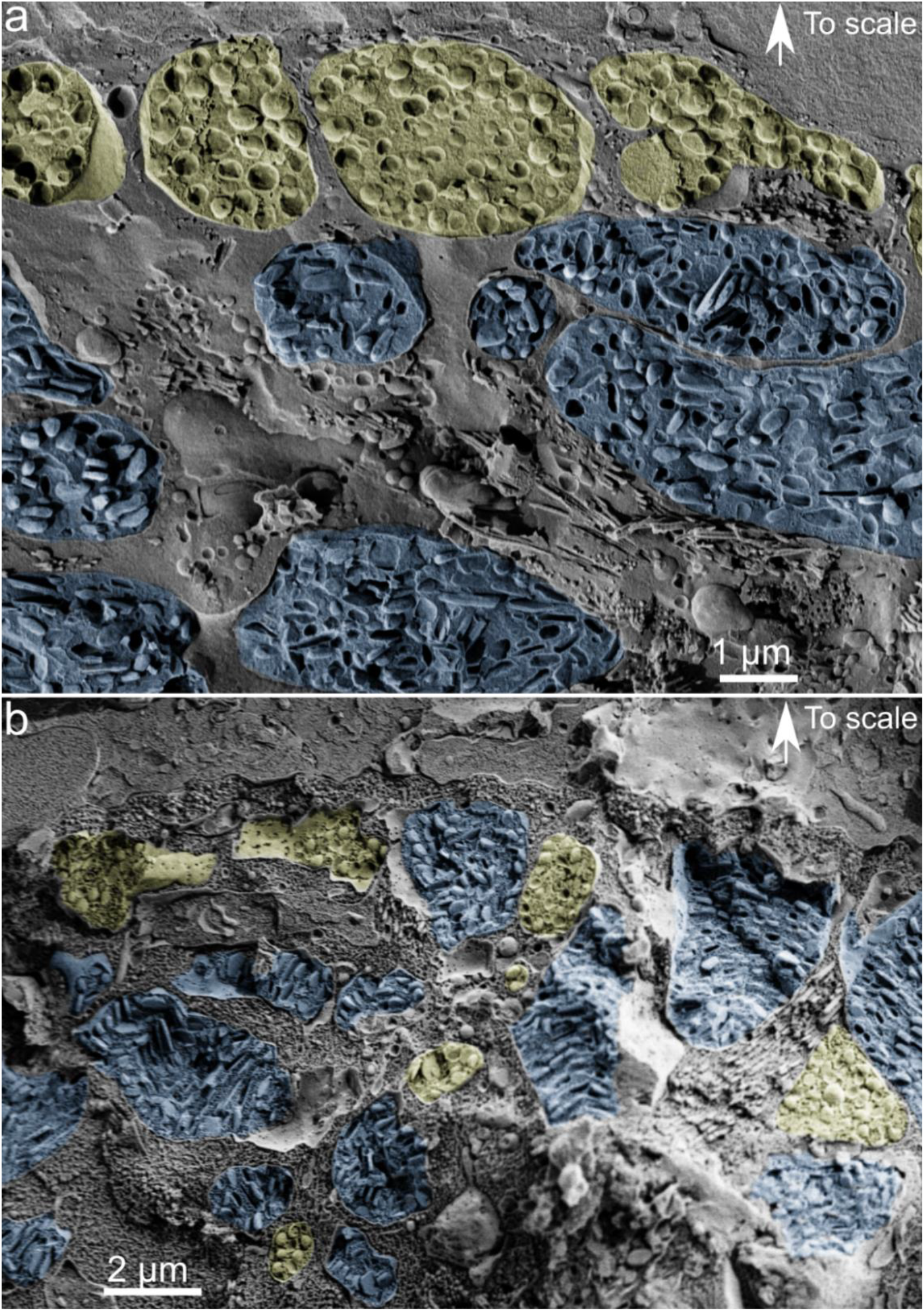
Cryo-SEM images of tail cross sections of *A. Beershebensis* from (**a**) hatchlings and (**b**) adults. Xanthophore cells are pseudo-coloured in yellow and iridophores in blue. The directions towards the scale are marked by white arrows.

**Extended Data Fig. 2.**
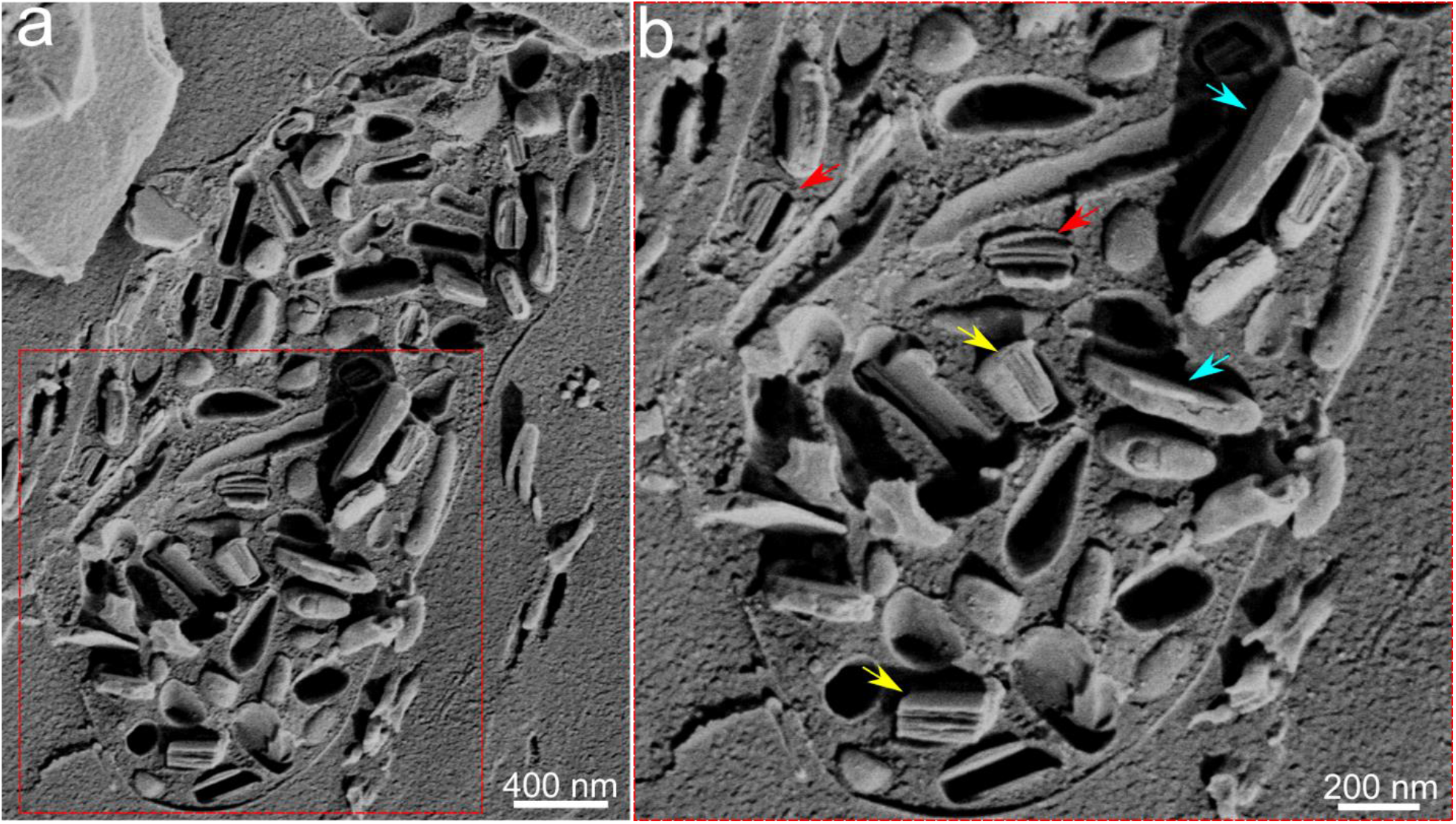
(**a**) Cryo-SEM images of iridophores from hatchling tails of *A. Beershebensis* with fractured vesicles showing partially formed guanine crystals. (**b**) Cryo-SEM images of fractured vesicles with guanine crystals at different formation stages. The coloured arrows represent a suggested sequence of formation steps: Red arrows: the most immature crystals, exhibiting distinctly separated platelets inside the ellipsoidal vesicles ^32, 33^. Yellow arrows: larger, more mature crystals containing more closely packed platelets. The vesicle membrane has been re-shaped by the crystal to adopt a faceted morphology. Cyan arrows: mature faceted in which individual platelets have coalesced.

**Extended Data Fig. 3.**
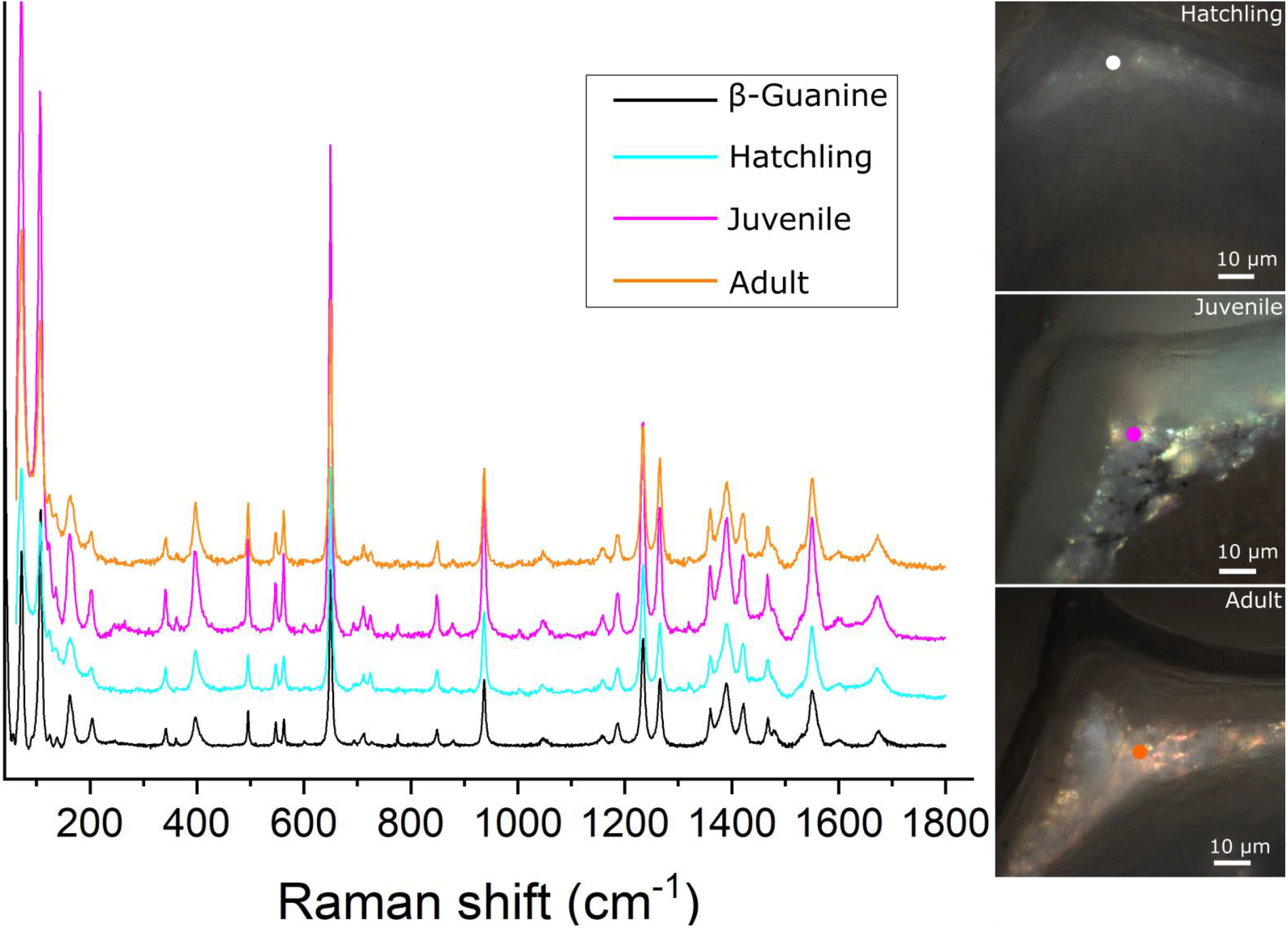
*In situ* Raman spectra obtained from the iridophore region in tail cross sections of *A. Beershebensis* from different developmental stages. The spectrum of β-guanine crystals is shown as a reference. The insets in the right column show corresponding optical micrographs of the tail cross sections. White, purple and orange dots in the micrographs denote the regions from which the spectra were obtained. The Raman spectra from each of the developmental stages is consistent with that of crystalline β-guanine.

**Extended Data Fig. 4.**
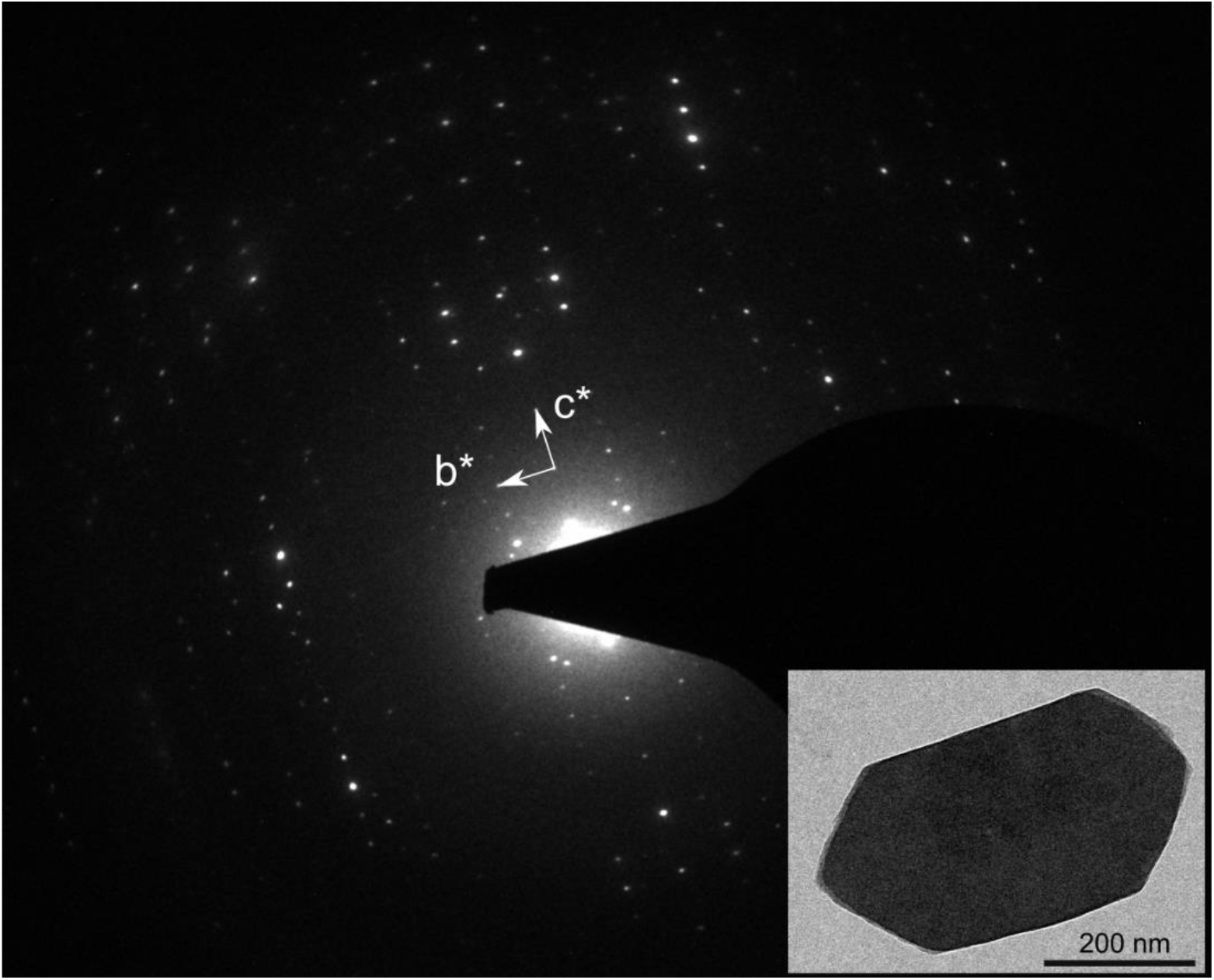
Electron diffraction pattern of a plate-like guanine crystal extracted from the iridophores of the tail of the adult *A. Beershebensis* (inset: real space TEM image of the corresponding crystal). The c* axis of the crystal can readily be identified from the measured periodicities. The length of the c* reciprocal axes = 0.054 Å^−1^ is consistent with the *d*_(001)_ spacing of β-guanine, 18.4 Å ^35^.

**Extended Data Fig. 5.**
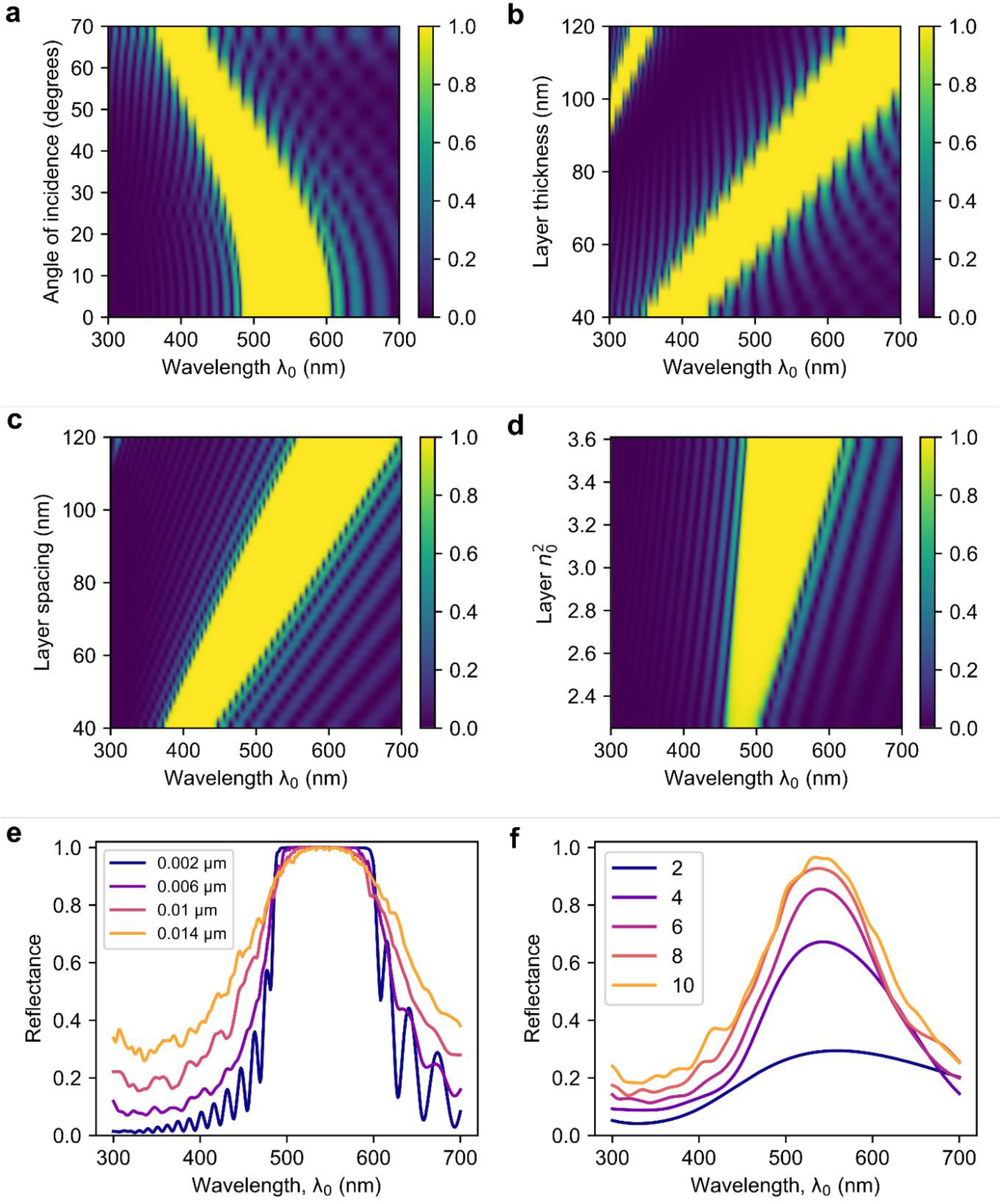
The variation of the reflectance spectra by different factors on a stack of 20 evenly spaced guanine/cytoplasm layers (layer thickness: 80 nm; layer spacing: 90 nm). (**a**) Variation in angle of incidence, indicating the blue shift of the high-reflectance peak as the angle of incidence increased. (**b**) Variation in the thickness of guanine layer. (**c**) Variation in the layer spacing. (**d**) Variation in the in-plane refractive index of the guanine layer at normal incidence, demonstrating the increasing refractive index will broaden the high-reflectance band. Both (**b**) and (**c**) indicate that changes to the structure of the layers result in a reduction of the length scale of the structural periodicity, which shifts the reflectance spectrum to blue. Structural correlations in the premature iridophores occur at the length scales smaller than the adult, as a result, any local layer like arrangement of the guanine platelets in premature iridophore would only enhances reflectance at the wavelengths shorter than in the adult. (**e**) Variation of reflectance spectrum on the stack of 20 spaced guanine/cytoplasm layers with magnitude of disorder. Upon introduction of equal amounts of disorder in the thickness and spacing of the layers (shown in the legend), the reflectance spectrum evolves from a sharp stop band to a rounded peak. The case of 0.014 µm disorder is close to the experimentally observed values. (**f**) Variation of the reflectance spectrum with number of layers (in the presence of disorder of 0.014 µm in both layer thickness and spacing). It can be observed that even a two layered structure is sufficient to exhibit a marked spectral feature in reflectance. In all calculations, disorder is quantified by the width (standard deviation) of the Gaussian distribution used to draw the values of parameters used in the calculations.

**Extended Data Fig. 6.**
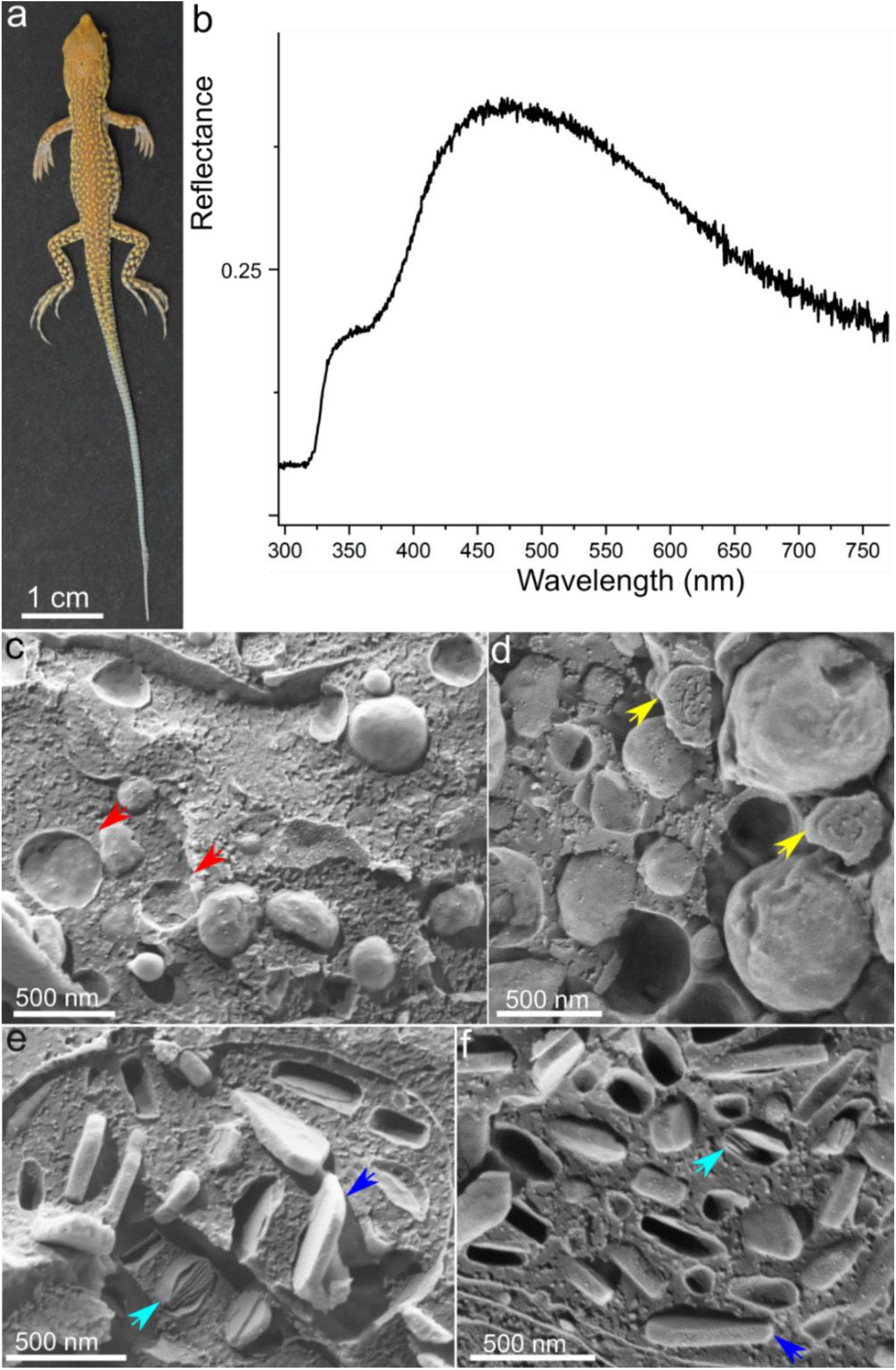
(**a**) Dorsal-lateral view of the hatchling blue tail *A. Scuttelatus* and (**b**) the corresponding reflectance spectrum. (**c, d**) Cryo-SEM images of fractured pterinosomes in the xanthophores. Empty pterinosomes are denoted by red arrows in C. Fibrous textures begin to emerge in some of the fractured pterinosomes (**d**, yellow arrows). (**e, f**) Guanine crystals in iridophores. Partially formed crystals are observed in the cells (**e, f**, cyan arrows). The blue arrow denotes mature guanine crystals.

